# A new single chain, genetically encoded biosensor for RhoB GTPase based on FRET, useful for live-cell imaging

**DOI:** 10.64898/2026.01.13.699355

**Authors:** Sandra Pagano, Louis Hodgson

## Abstract

RhoB is an atypical Rho GTPase whose function is tightly linked to its subcellular localization and membrane trafficking, reflecting its unique post-translational modifications and association with endosomal membranes in addition to the plasma membrane. Despite its implication in membrane trafficking and cytoskeletal regulation, tools to directly monitor RhoB activity in space and time have been lacking. Here, we describe the development and validation of a single-chain, genetically encoded Forster resonance energy transfer (FRET) biosensor that enables direct visualization of RhoB activity in living cells while preserving its native membrane-targeting determinants. The biosensor exhibits a large dynamic range and resolves spatially heterogeneous RhoB activity during leading-edge protrusion - retraction cycles in migrating mouse embryonic fibroblasts. To demonstrate the utility of this tool, we performed multiplex live-cell imaging with a previously developed near-infrared FRET biosensor for the exocytic Rho GTPase TC10. Quantitative morphodynamic and cross-correlation analyses reveal coordinated yet antagonistic spatiotemporal patterns of RhoB and TC10 activities at the leading edge and show that perturbation of TC10 regulation reorganizes their spatial coupling. Together, this work introduces a robust biosensor for RhoB and establishes a multiplex imaging framework to study the coordination of trafficking and signaling during cell migration.

## Introduction

RhoB is a member of the Rho subfamily within the Ras super-family of p21 small GTPases, sharing 71% and 78% amino-acid sequence identity with the canonical RhoA and another minor paralog RhoC, respectively (1). Rho GTPases function as molecular switches that cycle between an active GTP-bound and an inactive GDP-bound state to regulate a wide range of biological processes. Although the N-terminal regions of these three Rho GTPases are relatively conserved, their C-terminal regions diverge significantly, contributing to their distinct cellular localization and functions. Notably, RhoB contains a higher number of polar residues in its C-terminal hypervariable region and a distinct C-terminal CAAX box compared with RhoA and RhoC which together specify unique post-translational lipid modifications and subcellular targeting (2–4). Whereas RhoA and RhoC are exclusively geranylgeranylated, RhoB can additionally undergo palmitoylation as well as both geranylgeranylation and farnesylation (3, 5). Consequently, active RhoB is not confined to the plasma membrane but also localizes to endosomes and multivesicular bodies. (3).

Consistent with these properties, RhoB has been shown to play essential roles in multiple cellular processes, including migration (6–9), invasion (7–9), and proliferation (10, 11), through its involvement in cytoskeletal and adhesion dynamics, as well as membrane receptor recycling and intracellular trafficking of signaling molecules and membrane components. Despite these diverse functions, RhoB remains yet comparatively understudied (12–17). A deeper investigation of its activity and dynamics holds substantial promise for advancing our understanding of actin cytoskeleton regulation, cellular plasticity, and the intracellular trafficking pathways that support fundamental aspects of cell biology. Because the function of RhoB depends strongly on its subcellular compartmentalization and rapid cycling between active and inactive states, understanding its biology requires microscopy imaging tools capable of resolving its spatiotemporal activity in live cells, in the order of subcellular spatial and seconds temporal resolutions. In this context, the development of a dedicated single-chain, genetically encoded Förster resonance energy transfer (FRET) biosensor represents a powerful approach to overcome this limitation.

Genetically-encoded, single-chain fluorescent protein (FP)–based FRET biosensors represent a class of powerful microscopy imaging tools to visualize and monitor Rho GTPase activity in living cells with high spatial and temporal resolution(18–21). These biosensors consist of a FRET-donor and an -acceptor fluorescent proteins fused within a single polypeptide chain, together with the GTPase of interest and an effector-derived GTPase-binding domain (18, 19, 22–24). Such single-chain biosensors have been successfully developed for several Rho GTPases, including for the canonical members such as RhoA, Rac1, and Cdc42 (25–27), as well as other less extensively studied GTPases such as RhoC, Rac2, Rac3, and TC10 (28–31).

Here, we describe a new single-chain, genetically-encoded fluorescent protein–based FRET biosensor for RhoB GTPase. We retained full-length RhoB at the C-terminus of the FRET biosensor, including its C-terminal hypervariable region containing palmitoylation and prenylation sites that are essential for its characteristic localization at the plasma membrane and throughout the endosomal network (3, 5). By preserving these molecular determinants, the biosensor faithfully reports RhoB activity within its native subcellular compartments. The FRET-donor fluorescent protein is monomeric ECFP (32), and the -acceptor is a circularly permuted monomeric Citrine-YFP(33, 34), a configuration widely used in single-chain FRET biosensors to maximize dynamic range and sensitivity.

As a proof of concept, we monitored RhoB activity at the leading edge of migrating mouse embryonic fibroblasts (MEFs). To extend this analysis, we performed simultaneous imaging of our RhoB biosensor together with a previously reported near-infrared (NIR) FRET biosensor for the small GTPase TC10 (30), a key regulator of exocytosis (30, 35–38), enabling direct comparison of endocytic and exocytic signaling during cell migration. To mechanistically probe the coordination between these pathways, we perturbed p190RhoGAP, a GAP for TC10 with no known activity toward RhoB (30, 39), and examined whether modulation of TC10-dependent exocytic trafficking influences RhoB activity at the leading edge. Using this approach, we directly visualized the spatiotemporal regulation of RhoB in migrating cells and uncovered a coordinated, spatially organized pattern of RhoB and TC10 activities. Comparative analysis revealed complementary yet opposing activation dynamics, consistent with the engagement of interconnected but antagonistic signaling modules that regulate membrane remodeling and protrusive behavior during cell motility. Together, these results demonstrate the power of our RhoB biosensor to dissect the coordinated molecular mechanisms that drive leading-edge motion and, more broadly, cell motility dynamics.

## Results and discussion

We developed a genetically encoded single-chain FRET-based biosensor for RhoB GTPase. Building on our previously published RhoA biosensor (27), the new sensor is composed of full-length RhoB and a RhoB-specific binding domain derived from Protein kinase-N (PKN) RBD, positioned at the N- and C-termini, respectively (Fig. 1A). The terminal placement of RhoB preserves its C-terminal region, which is essential for RhoB subcellular localization and interaction with its regulators (1–3, 15, 40). The central region of the biosensor consists of two fluorescent proteins—monomeric ECFP and monomeric Citrine-YFP (41) which is circularly permuted at position 229 (42)—separated by a flexible linker of an optimized length (43) (Fig. 1A). The codon usage of the donor mECFP was synonymously modified to prevent spurious expression issues in living cells (44).

**Figure 1.**
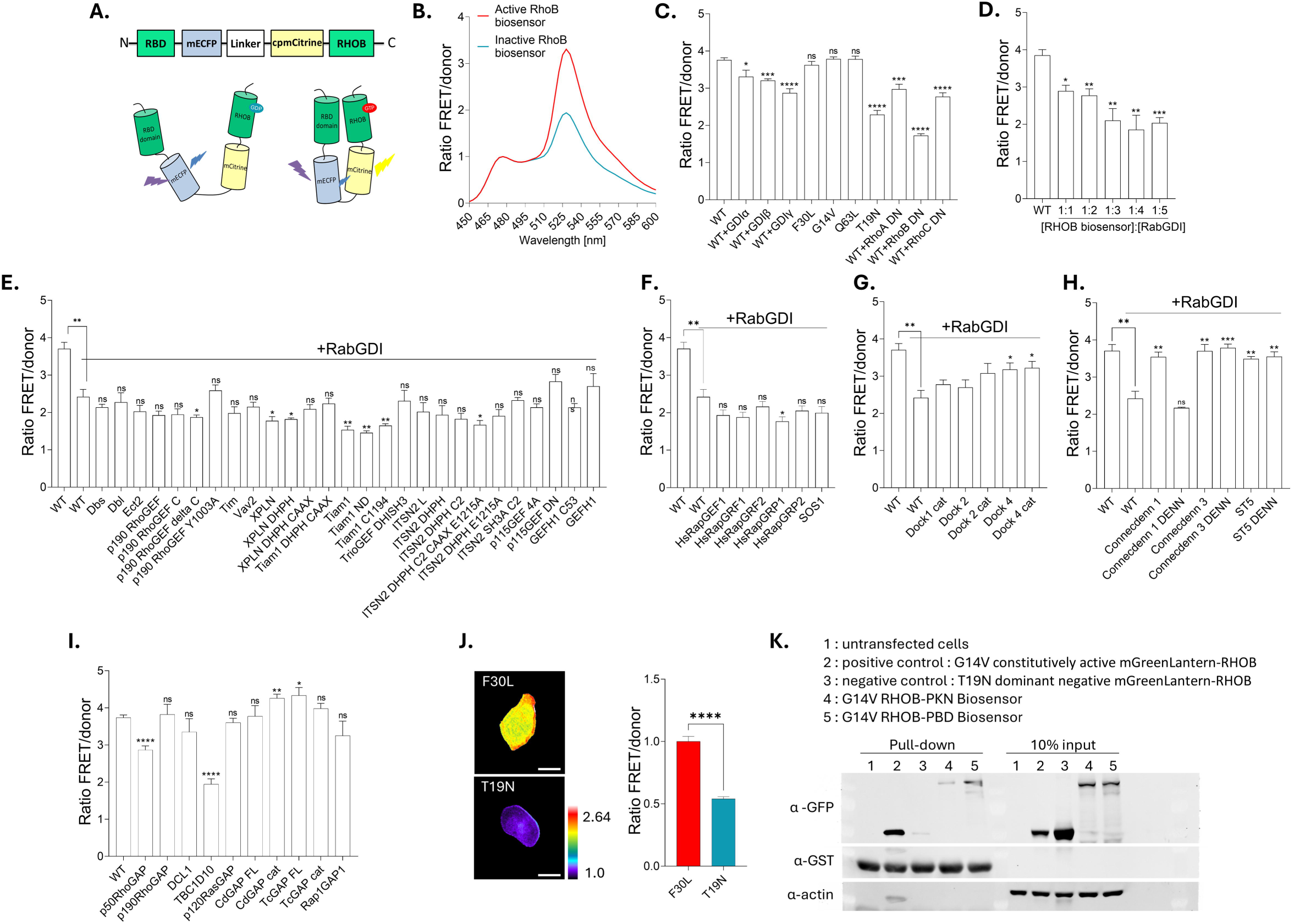
Design and validation of a CFP–YFP single-chain genetically encoded RhoB biosensor. **(A)** Schematic linear representation of the RhoB biosensor (top panel). From the N-terminus to the C-terminus, the biosensor is composed of the protein kinase N (PKN) Rho-binding domain (RBD), the donor fluorescent protein monomeric ECFP, a flexible linker, the acceptor circularly permuted monomeric Citrine, and full-length RhoB. The bottom panel illustrates the FRET mechanism, which results from the intramolecular interaction between GTP-bound RhoB and the PKN RBD. This interaction brings the donor and acceptor fluorescent proteins into close proximity upon the GDP-to-GTP switch, leading to increased FRET efficiency. **(B)** Representative normalized donor emission spectra of the RhoB biosensor harboring either a constitutively active mutation (G14V, red trace) or a dominant-negative mutation (T19N, blue trace). Data were obtained in HEK-293T cells following excitation at 433 nm, with normalization at the 474-nm emission peak. **(C)** Normalized FRET/CFP emission ratios of the wild-type (WT) RhoB biosensor alone or co-expressed with RhoGDI (α, β, or γ); of RhoB biosensor mutants (F30L, G14V, Q63L, T19N); and of the WT RhoB biosensor co-expressed with dominant-negative RhoA, RhoB, or RhoC in HEK-293T cells. Data are presented as mean ± SEM of 3-6 independent experiments. Statistical significance was assessed using an unpaired t-test. ns=non-significant, p > 0.05; *p < 0.05; ***p < 0.001; ****p<0.0001. **(D)** FRET/CFP emission ratios of the wild-type RhoB biosensor expressed alone or co-expressed with RabGDI. A titration was performed by varying the RhoB biosensor–to–RabGDI DNA ratio from 1:1 (200 ng of each plasmid) to 1:5 (200 ng RhoB biosensor plasmid and 1,000 ng RabGDI plasmid) in HEK-293T cells. Data are presented as mean ± SEM of 3 independent experiments. Statistical significance was assessed using an unpaired t-test. ns=non-significant, p > 0.05; *p < 0.05; **p<0.01; ***p < 0.001. **(E)** FRET/CFP emission ratios of the wild-type RhoB biosensor expressed alone or co-expressed with RabGDI and RhoGEFs in HEK-293T cells. Data are presented as mean ± SEM of 4 independent experiments. Statistical significance was assessed using an unpaired t-test. ns=non-significant, p > 0.05; *p < 0.05; **p<0.01. *n* = 4 independent experiments **(F)** FRET/CFP emission ratios of the wild-type RhoB biosensor expressed alone or co-expressed with RabGDI and RapGEFs in HEK-293T cells. Data are presented as mean ± SEM of 4 independent experiments. Statistical significance was assessed using an unpaired t-test. ns=non-significant, p > 0.05; *p < 0.05; **p<0.01. *n* = 4 independent experiments **(G)** FRET/CFP emission ratios of the wild-type RhoB biosensor expressed alone or co-expressed with RabGDI and DOCKGEFs in HEK-293T cells. Data are presented as mean ± SEM of 4 independent experiments. Statistical significance was assessed using an unpaired t-test. ns=non-significant, p > 0.05; *p < 0.05; **p<0.01. **(H)** FRET/CFP emission ratios of the wild-type RhoB biosensor expressed alone or co-expressed with RabGDI and RabGEFs in HEK-293T cells. Data are presented as mean ± SEM of 4 independent experiments. Statistical significance was assessed using an unpaired t-test. ns=non-significant, p > 0.05; *p < 0.05; **p<0.01; ***p < 0.001. **(I)** FRET/CFP emission ratios of the wild-type RhoB biosensor expressed alone or co-expressed with GAPs in HEK-293T cells. Data are presented as mean ± SEM of 5-7 independent experiments. Statistical significance was assessed using an unpaired t-test. ns=non-significant, p > 0.05; *p < 0.05; **p<0.01; ****p < 0.0001. **(J)** Representative ratiometric high-resolution microscopy images of transient over expression of the RhoB biosensor harboring either a constitutively active mutation (F30L) or a dominant-negative mutation (T19N) in MTLn3 cells (left panel), and quantification of the FRET/donor ratio (right panel). Scale bars=20 µm. Data are presented as mean ± SEM of 3 independent experiments, 31–37 cells per condition. Statistical significance was assessed using an unpaired t-test. ****p < 0.0001. **(K)** Western blot analysis of PKN-RBD pull-downs of monomeric GreenLantern–tagged RhoB mutants (G14V and T19N) used as assay controls, and of the constitutively active (G14V) RhoB biosensor containing either the original PKN Rho-binding domain (RBD) or a non-specific p21-binding domain (PBD), overexpressed in HEK-293 cells. Total cell lysates, GST, and actin were used as loading controls.

Because the size of the biosensor precluded *in vitro* purification, biosensor characterization by fluorescence measurements was performed in live, suspended HEK-293T cells (24, 45). Analysis of the fluorescence emission spectra of constitutively active (G14V) and dominant negative (T19N) mutants revealed an approximately 70% difference in FRET/donor ratio signal between the two states, demonstrating a large dynamic range of the biosensor (Fig. 1B). We further screened the activity of the RhoB WT biosensor in the presence of the different RhoGDI isoforms (GDIα, GDIβ, and GDIγ), which act as cytosolic chaperones that bind Rho GTPases and regulate their membrane association and activity (46–48) (Fig. 1C). Although the three RhoGDI reduced RhoB activity, GDIγ exerted the strongest inhibitory effect on the RhoB biosensor, consistent with previous reports identifying GDIγ as the only RhoGDI isoform documented to regulate RhoB (49, 50).

In parallel, we examined a panel of RhoB mutants, including a fast-cycling mutant (F30L), two constitutively active mutants (Q63L and G14V), and an inactive mutant (T19N) (Fig. 1C). No significant differences were detected between the active mutants and the WT biosensor, likely due to high expression levels of the constructs leading to saturation of the system and maximal activation of the WT biosensor (24, 27). As expected, the inactive mutant (T19N), for its part, exhibited reduced activity compared to WT and active mutants, consistent with the results shown in Fig. 1B. To demonstrate that the elevated activity of the WT biosensor likely stems from overexpression and aberrant activation by upstream regulators, namely guanine exchange factors (GEFs), we co-expressed non-fluorescent dominant-negative Rho GTPase mutants to titrate endogenous cellular GEFs (Fig. 1C). Indeed, co-expression of dominant-negative mutants of the three RhoGTPases decreased RhoB biosensor activity, with the most pronounced effect observed upon expression of dominant-negative RhoB, pointing to cellular endogenous GEF-mediated hyperactivation when WT biosensor was overexpressed. The next step was to test a panel of GEFs and GTPase-activating proteins (GAPs), which promote GTP loading and stimulate GTP hydrolysis, respectively, and assess their effects on the activity of the WT RhoB biosensor (Fig. 1E–I). This analysis is particularly relevant for RhoB, given the limited literature describing its upstream regulators. To facilitate the detection of GEF-dependent activation, we first reduced basal biosensor activity by co-expressing a GDI. Although GDIγ induced a measurable reduction in FRET/donor ratio, as shown by titration experiments (Supplementary Fig. 1), this effect remained relatively modest. Given the endosomal localization of RhoB and its close spatial and functional relationship with Rab GTPases, we next tested RabGDI. While RhoB biosensor activity was reduced by both RabGDI and GDIγ, RabGDI induced a stronger reduction than GDIγ. (Fig. 1D). Therefore, for subsequent screening experiments, we co-expressed the WT RhoB biosensor together with RabGDI and individual GEFs to assess the extent of rescue of RhoB biosensor activity. Co-expression of RhoGEFs (Fig. 1E) did not restore biosensor activity compared with RabGDI alone, consistent with an independent regulation of RhoB relative to other canonical Rho GTPases that are cytoskeleton-targeting. Interestingly, several RhoGEFs significantly decreased RhoB activity, suggesting an indirect inhibitory effect, potentially mediated through activation of other Rho GTPases. Screening of RapGEFs did not reveal any activation of RhoB (Fig. 1F). In contrast, DOCK family of GEFs targeting Rac1 and CDC42 (Fig. 1G) (51, 52), DOCK4—previously characterized as a Rac1 GEF—significantly increased RhoB activity. This observation is particularly interesting given the established role of RhoB in regulating Rac1 endosomal trafficking (6, 53). Strikingly, screening of RabGEFs resulted in robust restoration of WT RhoB biosensor activity across all tested conditions, highlighting a tight functional link between RhoB and the Rab GTPases, regulators of endosomal trafficking (Fig. 1H). Screening of GAPs further supported this observation, as RhoB activity was reduced upon expression of TBC1D10, a Rab-specific GAP, and p50RhoGAP, which has been shown to provide a functional link between Rab and Rho GTPases in the regulation of receptor-mediated endocytosis (54) (Fig. 1I).

We next examined differences in biosensor activity between constitutively active and dominant-negative RhoB mutants using high-resolution microscopy in the rat adenocarcinoma cell line MTLn3. Although subsequent experiments were performed in mouse embryonic fibroblasts (MEFs) stably and inducibly expressing the RhoB biosensor, these cells exhibit low transfection efficiency and are therefore not suitable for transfection-based overexpression of biosensor mutants. To overcome this limitation, MTLn3 cells were transiently transfected with RhoB biosensors harboring either the constitutively active F30L mutation or the dominant-negative T19N mutation. Biosensor activity was then quantified by wide-field fluorescence microscopy using ratiometric analysis (Fig. 1J). We observed an approximately twofold reduction in the FRET/donor ratio in the T19N mutant compared with the F30L mutant of RhoB, confirming the robust dynamic range of our biosensors. This experiment also showed the expected peri-membrane and cytoplasmic localization of the RhoB biosensor, pointing to the correct membrane and cytoplasmic partitioning due to the intact C-terminal region of the GTPase.

Next, we tested if the activated RhoB within the biosensor could compete against binding to endogenous downstream targets, which would result in strong overexpression artefacts. We produced a version of the constitutively activated G14V biosensor that contained a non-specific binding domain derived from p21-activated kinase 1 (PAK1-PBD), which is targeted by Rac1 and Cdc42 (55). PAK1-PBD was further rendered inert by inclusion of three additional point mutations (56–58). Glutathione S-transferase-fusion of PKN-RBD was used in excess to pull down the activated biosensor that contained an unmodified PKN-RBD or the inert PBD from the cell lysates. The pulldown assay confirmed that the excess exogenous binding domain can pull down the activated biosensor only when the internal binding domain was replaced with inert PAK1-PBD. This indicates that the internally accessible PKN-RBD and the RhoB within the biosensor backbone constitute the preferential interaction upon biosensor activation, minimizing overexpression artefacts (Fig. 1K).

After characterizing the biosensor under static conditions, the next step was to investigate its activity in live cells to demonstrate the biological relevance of the RhoB biosensor. Given that RhoB localizes to both the plasma membrane and endosomal compartments, we chose to examine RhoB activity at the leading edge of migrating cells (1, 13, 15, 17, 40, 59). To this end, we generated MEFs stably incorporating the RhoB biosensor under tetracycline inducible promoter using retroviral transduction followed by FACS sorting (27, 45). This approach ensures relatively low expression levels of the biosensors and a gated control of the biosensor expression population as needed for the microscopy imaging experiments. We quantified the biosensor expression levels in our stable cell line, relative to their endogenous counterparts by western blotting. The biosensor expression levels required to achieve sufficient signal to noise ratio in microscopy imaging was approximately 20% over the endogenous expression level of RhoB in these cells (Supplemental Fig. 3A). The required signal to noise ratio from the biosensor for imaging combined with the low endogenous expression levels of RhoB, resulted in modestly higher expression ratio over the endogenous protein levels than what we have used previously for this class of biosensors (27, 28, 30, 60). In biosensor-expressing cells, RhoB activity displays pronounced spatial heterogeneity, with elevated signals detected both at the cell periphery and within the cytoplasmic compartment. This distribution is consistent with the known subcellular localization of RhoB at the plasma membrane and endosomal structures (Fig. 2A–B). This representation illustrates the ability of the biosensor to resolve spatial variations in RhoB activity at the whole-cell level.

**Figure 2.**
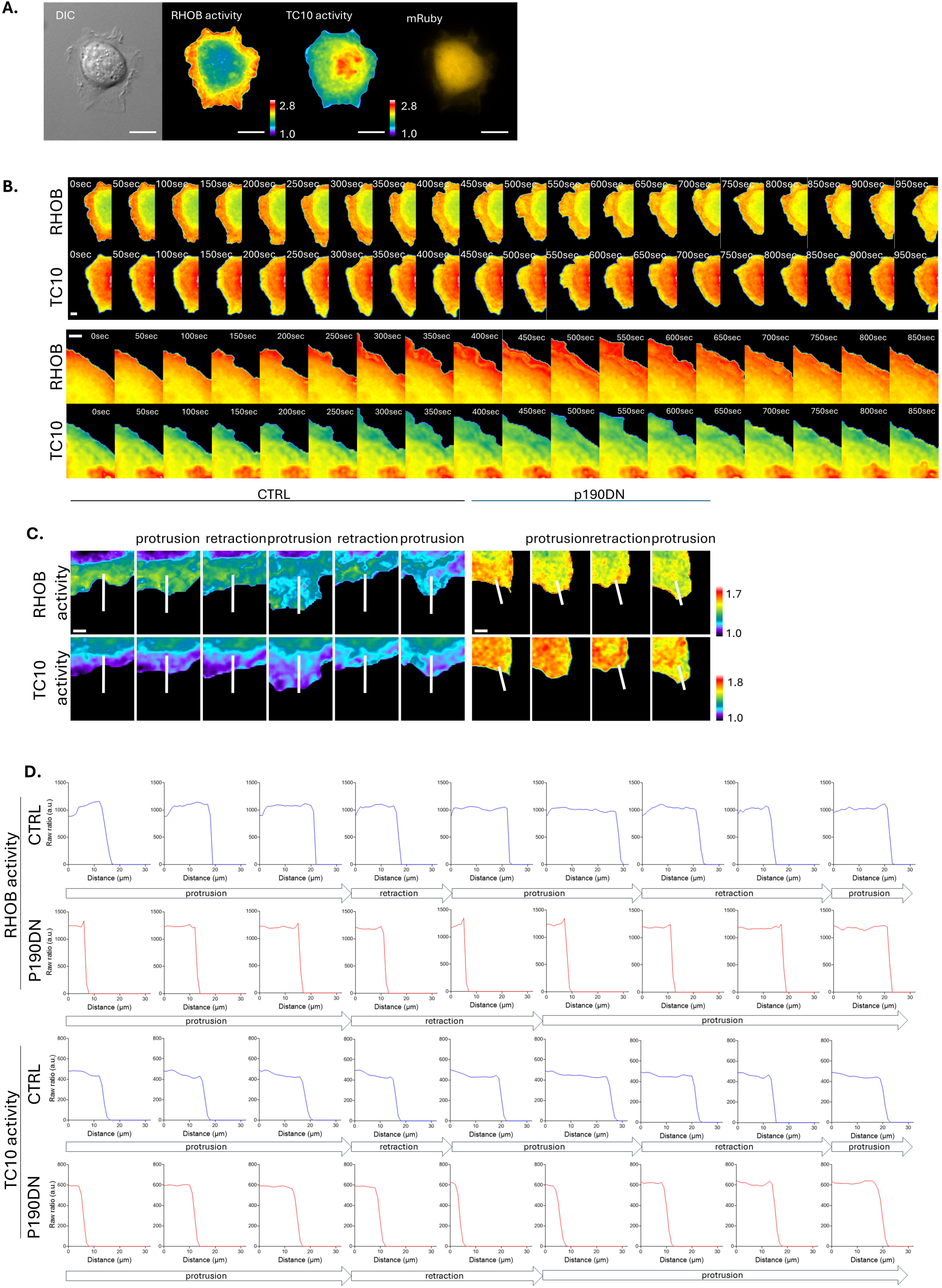
Spatial characterization of RhoB and TC10 biosensor activity in MEFs constitutively expressing both biosensors. **(A)** Representative high-resolution microscopy images of whole-cell MEFs shown in differential interference contrast (DIC), ratiometric images of stably expressed RhoB and TC10 biosensors, and transient expression of pTriEx-mRuby3 in control cells. Scale bars= 20 µm. **(B)** Representative high-resolution ratiometric time-lapse microscopy images of MEFs constitutively expressing RhoB and TC10 biosensors at the leading edge under control conditions. Scale bars: top panel= 10 µm, bottom panel = 5 µm. **(C)** Representative high-resolution ratiometric time-lapse microscopy images showing RhoB and TC10 biosensor activity during protrusion–retraction cycles in control and p190RhoGAP dominant-negative (p190DN)–transfected MEFs constitutively expressing both biosensors (top panels). The bottom panels show line-scan profiles of RhoB and TC10 activity along the line indicated in the first panel of the top row. The two upper line-scan panels depict RhoB activity in control (blue trace) and p190DN-transfected (red trace) MEFs, whereas the two lower line-scan panels depict TC10 activity in control (blue trace) and p190DN-transfected (red trace) MEFs. Scale bars= 5 µm.

To investigate the leading-edge dynamics, and specifically the balance between endocytic and exocytic compartments, we simultaneously monitored the exocytic network by stably co-expressing a previously published near-infrared TC10 biosensor from our laboratory (30), a key regulator of exocytosis (30, 39, 61). Like RhoB biosensor, TC10 biosensor was also virally transduced under inducible promoter and the expression levels were gated by FACS to achieve a minimum signal to noise ratio from the biosensor sufficient for microscopy imaging, at approximately 35% of the endogenous TC10 protein levels. We also assessed whether the induction of the biosensors affected endogenous protein expression levels. While no effect was observed on the endogenous TC10 protein level, endogenous RhoB level was slightly elevated when biosensors were expressed (Supplemental Fig. 3B), suggesting higher than endogenous expression levels of the RhoB biosensor could pose dominant negative effect leading to a minor compensatory increase in the endogenous RhoB expression levels.

In contrast to RhoB activity in cells, TC10 activity exhibits a markedly different spatial pattern (Fig.2A-B). Highest TC10 activity is concentrated in a perinuclear region, likely corresponding to the Golgi apparatus and/or the endoplasmic reticulum, compartments known to be essential for the maturation of TC10-positive exocytic vesicles (61). TC10 activity progressively decreases toward the plasma membrane as vesicles traffic away from their perinuclear sites of activation (Fig 2B). When focusing specifically on the leading edge during protrusion and retraction cycles, we observed an antagonistic pattern of RhoB and TC10 activities. RhoB activity was strongly enriched at the leading edge, whereas TC10 activity was relatively higher in more distal regions and progressively decreased toward the leading edge (Fig. 2B).

To further extend this analysis, we perturbed the balance between the endocytic and exocytic processes by expressing a dominant-negative (R1283A) form of p190RhoGAP (p190DN) (62). Previously, p190RhoGAP has been shown to target TC10 within the context of the leading-edge protrusion and cell migration in MDA-MB231 and HeLa cell lines (30, 39) but has not been reported to directly regulate RhoB. As noted previously, MEFs display low transfection efficiency. Therefore, p190DN was expressed as a construct coupled to the fluorescent protein mRuby3 (63), allowing visual identification of transfected cells. Control cells were transfected with the fluorescent protein alone, and only cells expressing the fluorescent marker were analyzed (Fig. 2A). Whole-cell–averaged analysis of RhoB and TC10 biosensor activity showed that p190DN expression did not significantly alter either biosensor signal (Supplementary Fig. 2A-B). Thus, p190RhoGAP, which is known to regulate its target GTPases including TC10 in a highly spatially and temporally restricted manner (30, 39), does not measurably alter TC10 activity when averaged across the entire cell. The lack of effect on RhoB activity is likewise expected, as RhoB has not been reported to be regulated directly by p190RhoGAP, and is consistent with our spectrofluorometric GAP screening data obtained in suspended HEK-293T cells (Fig. 1I). To more precisely analyze biosensor activity as a function of leading-edge dynamics, we performed line-scan analyses perpendicular to the cell edge under control conditions or upon expression of p190DN, and quantified biosensor activity during protrusion and retraction phases (Fig. 2C-D). Under control conditions, RhoB biosensor activity remained relatively uniform regardless of the distance from the leading edge, whereas p190DN expression was associated with a modest increase in RhoB activity at the leading edge, suggesting a possible enrichment of biosensor activity at the plasma membrane. Notably, under both control and p190DN conditions, no significant differences in RhoB activity were observed between protrusion and retraction phases (Fig. 2B). For the TC10 biosensor, control conditions revealed a slight decrease in activity at further distances away from the leading edge, consistent with the overall intracellular distribution of the biosensor activity. Interestingly, this spatial gradient was lost upon p190DN expression. Moreover, p190DN expression induced an increase in TC10 biosensor activity at the leading edge. These data are consistent with the previous findings that p190RhoGAP localized at the leading edge targets TC10 for GTP hydrolysis during cell protrusion and migration (39). As observed for RhoB, TC10 biosensor activity was not significantly modulated by protrusion–retraction cycles (Fig. 2B).

Following these high-level characterizations of leading-edge protrusions, we next applied morphodynamic mapping and cross-correlation analysis (64) to quantitatively relate leading-edge motion dynamics to the spatiotemporal activity patterns reported by the RhoB and TC10 biosensors at this site, a major hub of vesicular trafficking involving both endocytic and exocytic compartments. We restricted our study to non-migrating cells exhibiting robust protrusion–retraction fluctuations at the leading edge over the time course of a live cell experiment. Leading-edge movements were first analyzed using the temporal autocorrelation of the protrusion edge velocities, which revealed the characteristic periodic pattern of protrusion and retraction. This cyclic behavior and the characteristic periodicity (∼200s) were unaffected by p190DN compared to the control (Supplemental Fig. 4). Therefore, any observations of the biosensor behaviors described below reflect relative modulations of the activities of the GTPases under study rather than alterations in leading-edge protrusive behavior.

To further refine the analysis of biosensor dynamics, we examined the cross-correlation between the biosensor activities and leading-edge dynamics. Biosensor activity was quantified at increasing distances from the leading-edge using measurement windows of 6 pixels wide and 3 pixels deep, positioned from 0 to 30 pixels away from the edge, corresponding to 10 successive window strips (Fig. 3A) under control conditions or upon p190DN expression. For the RhoB biosensor, cross-correlation time-lag analysis revealed, under control conditions, a positive correlation between RhoB activity and protrusion, with RhoB activity lagging behind protrusion dynamics by 80 ± 44 s at the leading edge (Fig. 3B, black arrow). This positive correlation was maximal at the leading edge, progressively decreased up to approximately 12–15 pixels from the edge, and then increased again to reach a stable plateau in more distal windows (Fig. 3C). These observations indicate that the positive coupling between RhoB activity and protrusion dynamics is spatially restricted and not maintained uniformly with increasing distance from the leading edge. Upon expression of p190DN, no statistically significant differences were detected; however, a change in trend was apparent, characterized by a marked reduction of RhoB cross-correlation at the leading edge and a concomitant increase in more distal regions.

**Figure 3:**
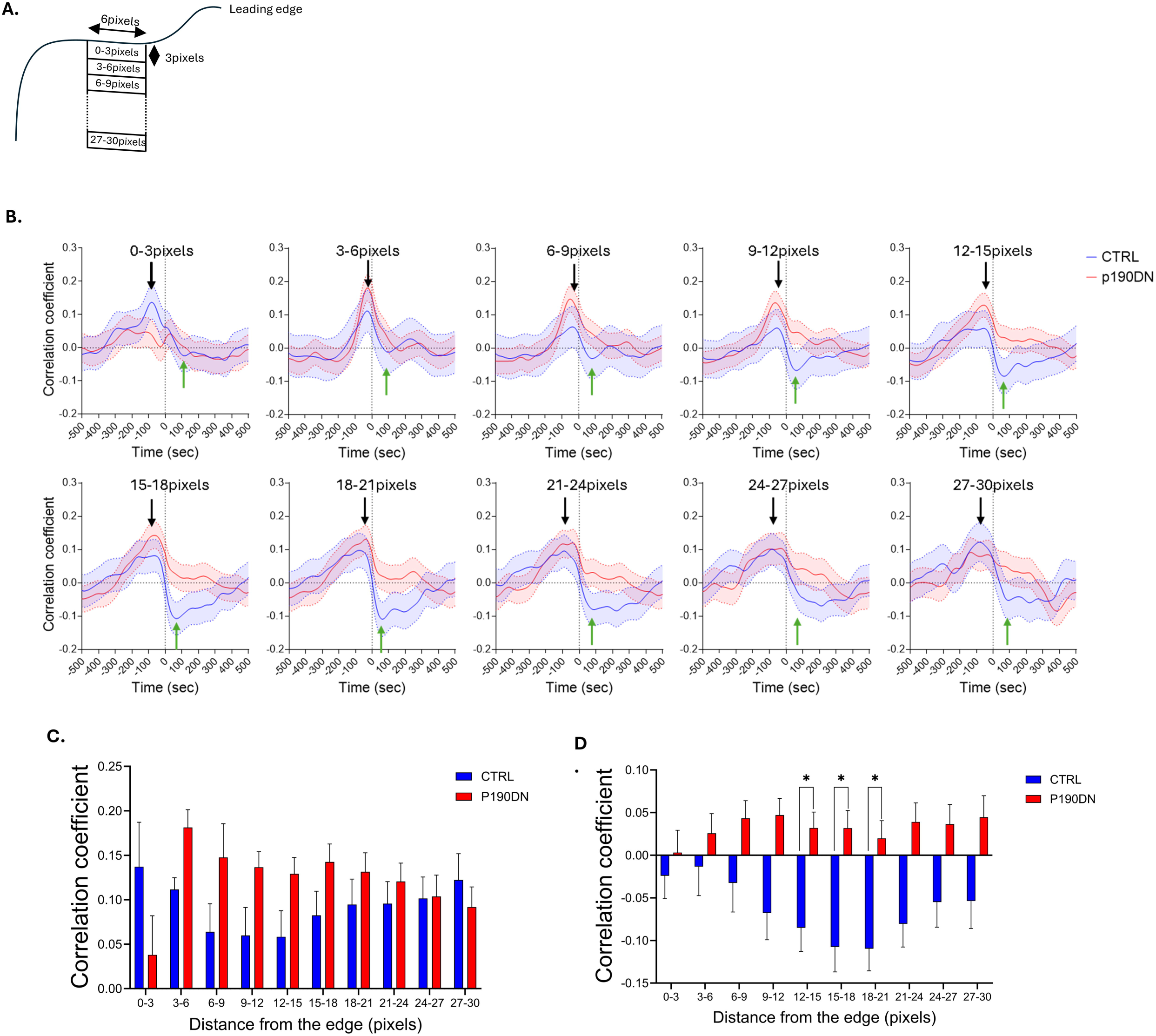
Morphodynamic analysis of RHOB activation dynamics during random protrusions in control and p190RhoGAP dominant-negative (p190DN)–transfected MEFs constitutively expressing RHOB and TC10 biosensors. RHOB activity was analyzed in MEFs co-expressing the RhoB and TC10 biosensors. The same cells were used for the quantification of both RhoB activity (Figure 3) and TC10 activity (Figure 4). **(A)** Schematic representation of the window strips designed for analysis and quantification of the protrusion–RhoB cross-correlation shown in panels (B)–(D). Ten window strips (6 × 3 pixels) were analyzed, spanning from the cell edge to more distal regions. Distances from the edge are defined as 0–3 pixels, 3–6 pixels, and so on. **(B)** RhoB activity and protrusion velocity cross-correlation in control and p190RhoGAP dominant-negative (p190DN)–transfected MEFs constitutively expressing both RhoB and TC10 biosensors. Correlation curves were computed from n = 535 individual windows across 12 cells from 4 independent experiments for the control condition, and n = 639 individual windows across 11 cells from 3 independent experiments for the p190DN condition. Data are presented as mean ± 95% CI. Black and green arrows indicate the locations used for quantification in panels (C) and (D), respectively. **(C)** RhoB activity and protrusion velocity cross-correlation at different distances from the cell edge, measured at the black arrow position indicated in panel **(B)**, in control (blue trace) and p190RhoGAP dominant-negative (p190DN)–transfected (red trace) MEFs constitutively expressing both RhoB and TC10 biosensors. Correlation curves were computed from n = 535 individual windows across 12 cells from 4 independent experiments for the control condition, and n = 639 individual windows across 11 cells from 3 independent experiments for the p190DN condition. Data are presented as mean ± 95% CI. Statistical significance was assessed using an unpaired t-test. **(D)** RhoB activity and protrusion velocity cross-correlation at different distances from the cell edge, measured at the green arrow position indicated in panel **(B)**, in control (blue) and p190RhoGAP dominant-negative (p190DN)–transfected (red) MEFs constitutively expressing both RhoB and TC10 biosensors. Correlation curves were computed from n = 535 individual windows across 12 cells from 4 independent experiments for the control condition, and n = 639 individual windows across 11 cells from 3 independent experiments for the p190DN condition. Data are presented as mean ± 95% CI. Statistical significance was assessed using an unpaired t-test. *p < 0.05.

In contrast, when focusing on positive time lags relative to edge protrusion, RhoB exhibited a distinct behavior (Fig. 3B, green arrow–C). Under control conditions, we observed the progressive emergence of a negative correlation between protrusion and RhoB activity with increasing distance from the leading edge. This relationship was associated with a temporal delay of protrusion dynamics relative to RhoB activity leading protrusion initiation by 68 ± 51 s at 12-15 pixels from the edge, indicating that RhoB activity precedes protrusive events and is negatively correlated with protrusion in these more distal regions (Fig. 3D). Upon expression of p190DN, this negative cross-correlation was abrogated (Fig. 3D). These results indicate that p190RhoGAP is required for the spatial and temporal compartmentalization of RHOB activity. Although p190RhoGAP has not been previously reported to directly regulate RhoB, our data suggests that p190RhoGAP influences the coupling of the edge movement to the sequential, spatiotemporal regulation of RhoB activity. Notably, the temporal offset between the positive and negative correlations was shorter than the duration of a protrusion event, indicating that these opposing relationships occur within the same protrusion–retraction cycle. Together, these data support a sequential pattern in which RhoB activity transitions from a positive to a negative correlation with protrusion dynamic during a single protrusion cycle. Thus, p190RhoGAP-dependent spatiotemporal modulation of RhoB activity appears to be important during the dynamics of leading-edge protrusion and retraction. This may be explained by the fact that RhoB operates at the intersection of endosomal trafficking and intracellular signaling. Through its ability to regulate the trafficking of other small GTPases, RhoB may influence their spatial activation and inactivation cycles and thereby modulate cytoskeletal regulators involved in leading-edge dynamics (53, 65, 66). In parallel, by controlling growth factor receptor recycling, RhoB may contribute to the spatial and temporal coordination of protrusive responses to extracellular cues (4, 67). Together, these complementary mechanisms could underlie the role of RhoB in shaping protrusion–retraction dynamics at the leading edge.

We performed the same analysis for TC10 activity in relation to protrusion dynamics (Fig. 4A–B). Under control conditions, a strong negative cross-correlation between protrusion and TC10 activity was observed at the leading edge with no detectable time lag (Fig. 4A). This negative correlation progressively decreased with increasing distance from the edge and was not detected in more distal regions, indicating that the relationship between TC10 activity and protrusive dynamics is spatially restricted to the leading edge. These results show that TC10 activity is inversely related to protrusion specifically at the leading edge, consistent with previous observations that TC10 GTP hydrolysis at this site is required for efficient migration by promoting fusion of exocytic vesicles through exocyst complex docking (30, 39). This spatially confined inverse relationship further suggests that TC10 activity is preferentially elevated during leading-edge retraction. Given that TC10-driven exocytosis supplies lipids to the plasma membrane (38, 68), such fluctuations in TC10 activity may contribute to protrusion dynamics through cycles of membrane expansion and stabilization. Expression of p190DN strongly attenuated the negative cross-correlation at the leading edge, while the overall pattern was otherwise preserved, suggesting that p190RhoGAP-dependent TC10 inactivation is required to properly establish the spatiotemporal relationship between TC10 activity and protrusion dynamics (Fig. 4B).

**Figure 4:**
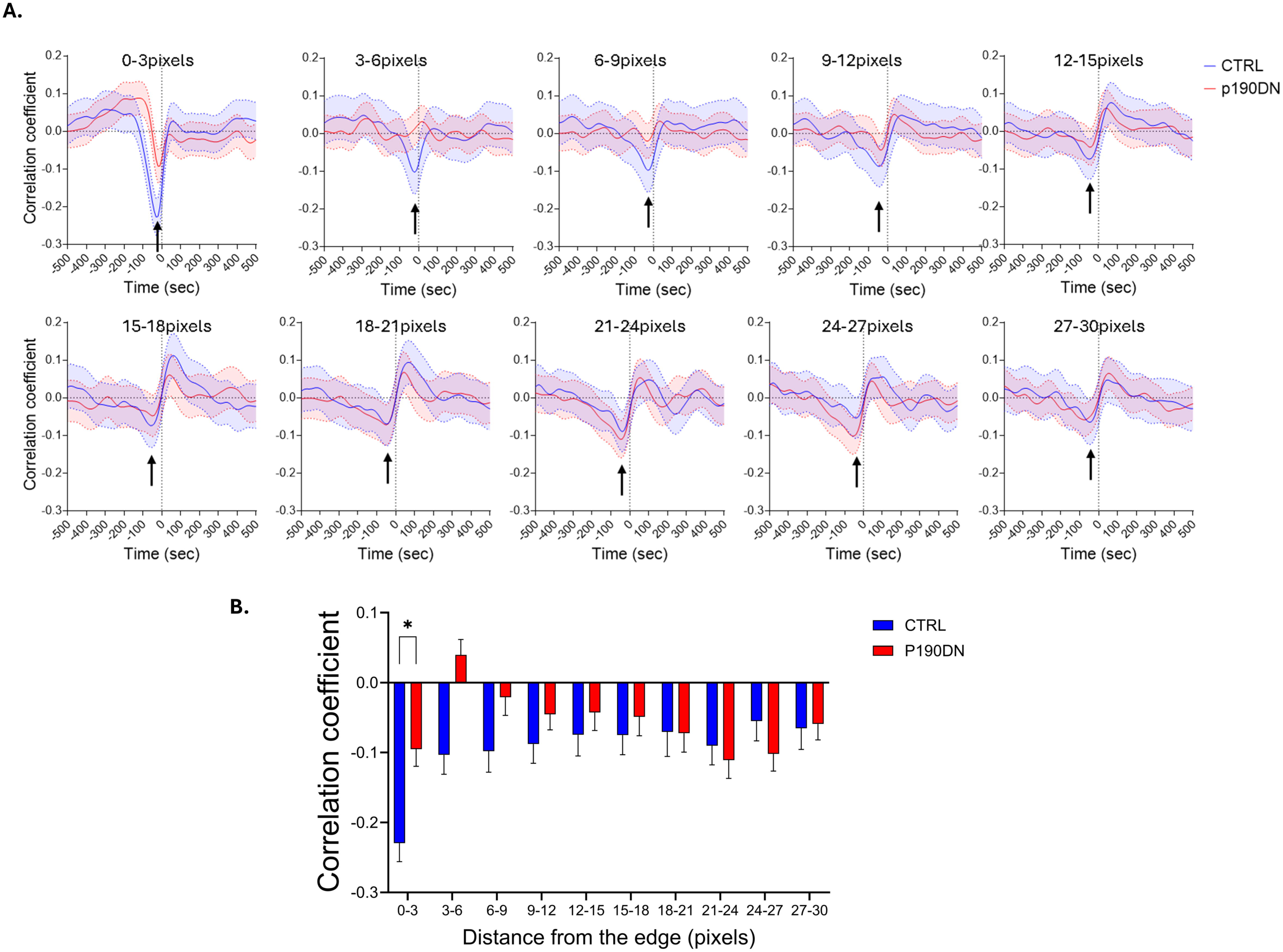
Morphodynamic analysis of TC10 activation dynamics during random protrusions in control and p190RhoGAP dominant-negative (p190DN)–transfected MEFs constitutively expressing RHOB and TC10 biosensors. RHOB activity was analyzed in MEFs co-expressing the RhoB and TC10 biosensors. The same cells were used for the quantification of both RhoB activity (Figure 3) and TC10 activity (Figure 4). **(A)** TC10 activity and protrusion velocity cross-correlation in control and p190RhoGAP dominant-negative (p190DN)–transfected MEFs constitutively expressing both RhoB and TC10 biosensors. Correlation curves were computed from n = 535 individual windows across 12 cells from 4 independent experiments for the control condition, and n = 639 individual windows across 11 cells from 3 independent experiments for the p190DN condition. Data are presented as mean ± 95% CI. **(B)** RhoB activity and protrusion velocity cross-correlation at different distances from the cell edge, measured at the black arrow position indicated in panel **(A)**, in control (blue trace) and p190RhoGAP dominant-negative (p190DN)–transfected (red trace) MEFs constitutively expressing both RhoB and TC10 biosensors. Correlation curves were computed from n = 535 individual windows across 12 cells from 4 independent experiments for the control condition, and n = 639 individual windows across 11 cells from 3 independent experiments for the p190DN condition. Data are presented as mean ± 95% CI. Statistical significance was assessed using an unpaired t-test. *p < 0.05.

Because the distribution of the cross-correlation functions for the protrusion vs RhoB or TC10 is complex, we next sought to examine the direct cross-correlation between RhoB and TC10 activities, made possible by our direct multiplex biosensor imaging modality (Fig. 5A–B). We observed no correlation between RhoB and TC10 activities at the leading edge (0–3 pixels window), as expected from their relative behaviors at the leading edge when cross-correlated against the protrusion velocities. Interestingly, a strong negative cross-correlation emerged in regions located 3–21 pixels away from the edge, which progressively weakened and ultimately disappeared at more distal positions (Fig. 5A). Thus, while the individual measurements and analysis of RhoB and TC10 activities appear spatially complex at the leading edge, the pronounced and clear negative cross-correlation observed further from the edge supports a direct, antagonistic relationship between these two RhoGTPases and, indirectly, between the cellular processes they regulate—endocytosis and exocytosis, respectively. Importantly, this direct negative cross-correlation between RhoB and TC10 activities was fully abrogated upon expression of p190DN, suggesting that p190RhoGAP plays an important role in the spatiotemporal regulation, compartmentalization and coordination of RhoB and TC10 activities at the leading edge (Fig. 5B).

**Figure 5:**
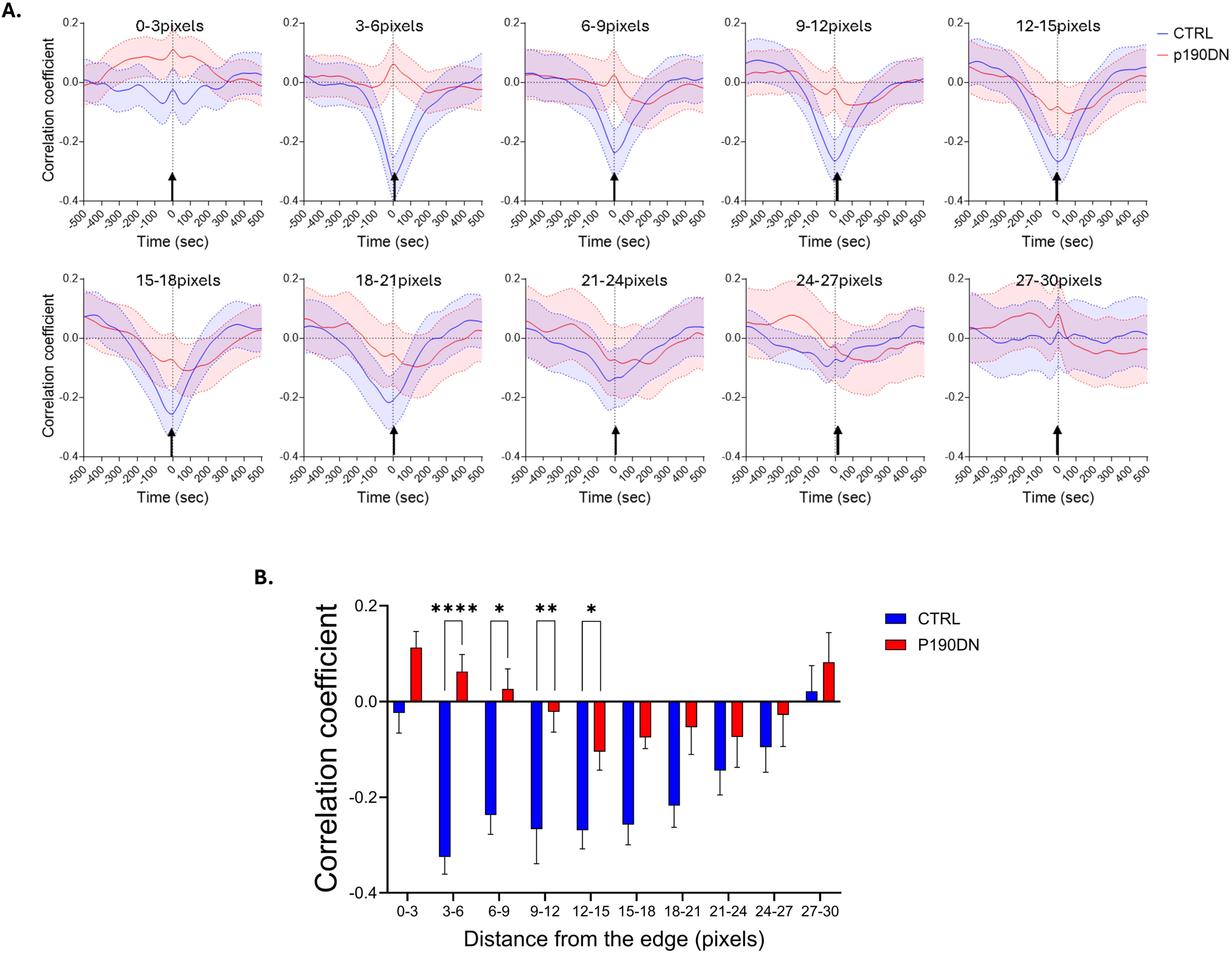
Morphodynamic analysis of RHOB and TC10 activation dynamics during random protrusions in control and p190RhoGAP dominant-negative (p190DN)–transfected MEFs constitutively expressing RHOB and TC10 biosensors. **(A)** Correlation of RHOB activity and TC10 activity monitored in the same cell in control and p190RhoGAP dominant-negative (p190DN)–transfected MEFs constitutively expressing both RhoB and TC10 biosensors. Correlation curves were computed from n = 535 individual windows across 12 cells from 4 independent experiments for the control condition, and n = 639 individual windows across 11 cells from 3 independent experiments for the p190DN condition. Data are presented as mean ± 95% CI. **(B)** RhoB activity and TC10 activity cross-correlation at different distances from the cell edge, measured at the black arrow position indicated in panel **(A)**, in control (blue trace) and p190RhoGAP dominant-negative (p190DN)–transfected (red trace) MEFs constitutively expressing both RhoB and TC10 biosensors. Correlation curves were computed from n = 535 individual windows across 12 cells from 4 independent experiments for the control condition, and n = 639 individual windows across 11 cells from 3 independent experiments for the p190DN condition. Data are presented as mean ± 95% CI. Statistical significance was assessed using an unpaired t-test. *p < 0.05; **p<0.01; ****p<0.0001.

## Methods

### Cell culture

MTLn3 cells (rat adenocarcinoma) (69) were cultured in Minimum Essential Medium (MEM, Corning, Corning, NY, USA) supplemented with 5% fetal bovine serum (FBS), 1% glutamine, and 100 I.U. penicillin and 100□µg/mL streptomycin (Invitrogen, Carlsbad, CA, USA), as previously described (70). MEF and HEK-293T cells were cultured in Dulbecco’s modified Eagle medium (DMEM, Corning) supplemented with 10% FBS, 1% glutamine, and penicillin/streptomycin, as previously described (71).

### Transfection

For MEF cell line, plasmid transfections were performed in OptiMEM, using TurboFect transfection reagent (Thermo Scientific). Cells were plated at 5□×□10^4^ cells/well in a 6-well plate and incubated overnight prior to transfection. Following the manufacturer’s protocols, 4□µg of total DNA was transfected into each well of a 6-well plate. Cells were treated with the transfection mixture overnight. For MTLn3 cell line, transfections were performed using Lipofectamine 2000 (Invitrogen). Cells were plated at 2□×□10^5^ cells/well of a 6 well plate and allowed to adhere and grow overnight. The following day 2 µg of DNA was added to 250 µl of OptiMEM and 4 µl of Lipofectamine 2000 was added to a separate aliquot of 250 µl of OptiMEM and incubated at room temperature for 5 minutes. The Lipofectamine solution was added to the DNA mixture and incubated for 20 minutes at room temperature. Cells were washed once with PBS, and 500 µl of OptiMEM was added to the well. Transfection mixture was added and the media was replaced with normal culture medium 45 minutes later and allowed to grow overnight.

### Biosensor construction and production of stable-inducible RhoB MEF cells

A FRET biosensor for RhoB was constructed based on the previously published RhoA, single-chain, genetically encoded biosensor backbone system (27). Briefly, WT and mutant human RhoB GTPase sequences were PCR-amplified using the primer pair: 5’- GCTTAATTAGTTGCAAGAATTCATGGCGGCCATCCGCAAGAAGCTGGT-3’ and 5’- GCAAATATGAATTCTTACTCGAGTCATAGCACCTTGCAGCAGTTGATGCA-3’ and restriction digested with EcoRI and XhoI. The digested fragments were ligated into the pTriEX-4 vector containing the RhoA FRET biosensor backbone at the EcoRI/XhoI sites to exchange the RhoA GTPase sequence for the RhoB GTPase fragments. The binding domain was replaced from the original Rhotekin-RBD to Protein kinase N-RBD (1-100 amino acids) (72), by PCR amplification using primer pair: 5’- GCAAATATGAATTCTTACCATGGCCAGCGACGCCGTGCAGAGTGA-3’ and 5’- GCTAATGTAACAAGTATGGATCCAAGCACCACGTGGGCGTGCAGCTCCT-3’ followed by digestion with NcoI/BamHI and ligation into the biosensor backbone. The donor fluorescent protein was replaced from the original ECFP (with intact dimerization interface containing A206) to the synonymous codon modified (44) monomeric ECFP (incorporating A206K mutation) (73) through subcloning from our second generation monomeric RhoA biosensor (74), at BamHI/HindIII sites. Circularly permutated monomeric Citrine-YFP was generated using 2-step PCR amplification, with monomeric Citrine (41) as the template and using the following primer sets: 5’- GGTATTAATTAATATGCGGCCGCTATGATCACTCTCGGCATGGACGAGC-3’, 5’- GCCACCGCTGCCACCCTTGTACAGCTCGTCCATGCCGAGAGTGATCAT-3’, 5’- GCTGTACAAGGGTGGCAGCGGTGGCATGGTGAGCAAGGGCGAGGAGCTGT-3’, and 5’- CCATATTAATATATGAATTCCTTCCCGGCGGCGGTCACGAACTCCAGCA-3’, digested and ligated into the biosensor backbone at NotI/EcoRI sites. To generate the retroviral vector containing the biosensor in the tet-inducible system, the pRetro-X-DEST vector system (Clontech, Mountainview, CA, USA) was used. The pTriEX-RhoB biosensor was digested using NcoI and XhoI to extract the RhoB biosensor as a full-length cassette, which was then ligated into the pENTR-4 vector (Invitrogen) at NcoI/XhoI sites. The pENTR-RhoB biosensor was processed using Gateway cloning technique, together with the pRetro-X-Puro-DEST vector, using LR Clonase II (Invitrogen), following the manufacturer’s protocols. The pRetro-X-Puro-RhoB biosensor was used to produce retrovirus for infecting MEF cells to produce stable/inducible tet-OFF biosensor cell line, as previously described (75, 76). The base pair sequence information for RhoB biosensor is shown in Supplementary Data 1.

### Fluorometry

Biosensor response characterization was performed in LinXE cells (a HEK293T-derived cell line) (77, 78) by transient expression of wild-type or mutant biosensor constructs, with or without upstream regulatory proteins, as described previously (27). LinXE cells were plated overnight on poly-L-lysine–coated 6-well plates (Sigma) at a density of 9 × 10^5^ cells per well and transfected the following day using Polyethylenimine following the optimized procedures (79). Biosensors were co-transfected with regulatory proteins at the following plasmid ratios: 1:4 for co-expression with GDI, DN, GAPs or GEFs. Forty-eight hours post-transfection, cells were washed with PBS, briefly trypsinized, and resuspended in cold PBS. Live-cell suspensions were transferred to a quartz cuvette, and fluorescence emission spectra from 450 to 600 nm were acquired using a spectrofluorometer (Fluorolog-3 MF2; Horiba Jobin Yvon). Samples were excited at 433 nm. Background spectra obtained from cells transfected with empty vector were used to correct for light scatter and cellular autofluorescence. Corrected spectra were normalized to the peak mECFP emission intensity at 475 nm to generate ratiometric emission profiles.

### Pull down assay

Pull-down assays were performed using purified PKN-RBD–conjugated agarose beads, as previously described (58). Glutathione (GSH)-agarose beads (Sigma-Aldrich) were prepared by resuspending 100 mg of beads in sterile water, incubating for 1 hour at 4 °C, and washing three times with water followed by two washes in resuspension buffer (50 mM Tris, pH 8.0, 40 mM EDTA, 25% sucrose). The beads were finally resuspended in 2 ml of resuspension buffer. To generate GST–PKN-RBD, the PKN Rho-binding domain (amino acids 1–100) was amplified by PCR and subcloned into the pGEX-4T1 vector (Cytiva) using BamHI and XhoI restriction sites. The resulting construct was transformed into BL21(DE3) competent bacteria (Agilent Technologies). Bacterial cultures were grown at 37 °C with shaking (225 rpm) to an OD□□□ of ∼1.0, and protein expression was induced with 0.2 mM IPTG. Cultures were immediately shifted to 25 °C and incubated overnight.

Cells were harvested and resuspended in resuspension buffer supplemented with 1 mM PMSF, protease inhibitor cocktail (Sigma-Aldrich), and 2 mM β-mercaptoethanol, followed by incubation at 4°C for 20 minutes. Detergent buffer (50 mM Tris, pH 8.0, 100 mM MgCl₂, 0.2 % Triton X-100) was added, and lysates were incubated for an additional 10 minutes at 4 °C. Cells were lysed by ultrasonication (4 × 45 seconds cycles on ice) and clarified by centrifugation at 22,000 rcf for 45 minutes at 4 °C. The supernatant was incubated with prepared GSH-agarose beads for 1 hour at 4 °C with rotation. Beads were washed four times with wash buffer (50 mM Tris, pH 7.6, 50 mM NaCl, 5 mM MgCl₂) and resuspended in 50 % glycerol/wash buffer. Aliquots (100 µl) were flash-frozen in liquid nitrogen and stored at −80 °C until use.

For pull-down experiments, LinXE cells were transfected as indicated and lysed in RBD pull-down lysis buffer (50 mM Tris, pH 7.4, 500 mM NaCl, 50 mM MgCl₂, 1 % NP-40). Lysates were sonicated, clarified by centrifugation (22,000 rcf, 15 minutes, 4 °C), and an input fraction was retained. The remaining lysates were incubated with PKN-RBD–agarose beads for 1 hour at 4 °C, washed four times in lysis buffer, resuspended in SDS sample buffer, and analyzed by western blotting. RhoB biosensor or fluorescently tagged RhoB proteins were detected using anti-GFP antibody (mouse; Roche, clones 7.1 and 13.1).

### Microscopy imaging

MEF cells stably expressing the indicated biosensors were plated on 25-mm round #1.5 glass coverslips (Warner Instruments) coated with fibronectin (Sigma; 10 μg /ml in PBS, 1 hour at room temperature) at a density of 5 × 10□ cells per well. Cells were maintained in standard growth medium supplemented with 25 μM biliverdin (BV; Sigma) on the day of the experiment and imaged 3 h after plating. Live-cell imaging was performed at 37 °C in a closed imaging chamber using Fluororbrite DMEM without phenol red (Invitrogen), sparged with argon gas to reduce dissolved oxygen and supplemented with 3 % fetal bovine serum, Oxyfluor reagent (1:100; Oxyrase Inc.), and 10 mM dl-lactate (Sigma) (80). Imaging medium did not contain exogenous BV.

Widefield FRET imaging was carried out using a custom Olympus IX83-ZDC2 microscope optimized for ratiometric biosensor imaging(71). Image acquisition was controlled using Visiview v7.0.0.8 (Visitron Systems GmBH). Images were acquired through a 40× 1.3 NA oil-immersion objective (Olympus UIS DIC) with 2 × 2 camera binning. Simultaneous acquisition of mECFP and mCitrine emissions was achieved using two synchronized PrimeBSI-Express sCMOS cameras (Photometrics) mounted on a 4-way beamsplitter (Cairn) attached onto the left-side emission port of the microscope. The beamsplitter was equipped with appropriate dichroics and emission filters (Chroma Technology) (71). Two additional PrimeBSI sCMOS cameras (Photometrics) were mounted on this beamsplitter at the terminal position to simultaneously acquire miRFP670/miRFP720 FRET channels for the NIR FRETbiosensor. Inclusion of a filterwheel (Ludl Electronic Products) in one of the terminal cameras enabled acquisition of mRuby3 as the transfection marker.

All image channels were registered prior to ratiometric calculations using pixel-by-pixel alignment based on *a priori* calibration and non-linear coordinate transformation(80, 81). Image processing was performed using MetaMorph and MATLAB (MathWorks) and included flat-field correction, background and camera noise subtraction, threshold masking, ratiometric calculations, photobleaching correction, and spatial registration, as previously described (81).

### Morphodynamic-mapping and cross-correlation analysis

Morphodynamic mapping and cross-correlation analyses were performed as described previously (64). Briefly, cell edge motion was tracked from time-lapse image series, and measurement window segments (3 × 6 pixels) were constructed along the leading edge to quantify biosensor activities and edge velocity during complete protrusion–retraction cycles. This window size was previously determined to be diffusion-limited under the imaging conditions used, allowing each segment to be treated as an independent sampling entity. Measurement windows were progressively displaced from the leading edge in 3 pixel increments to assess spatial coupling of biosensor activities. Temporal relationships between biosensor readouts were quantified using cross-correlation analysis (*xcov*, MATLAB), with Pearson’s correlation coefficient used to assess the coupling strength. Statistical confidence intervals (95%) were determined by bootstrapping (2000 iterations) of spline-smoothed correlation functions. A total of 535 window segments from 12 cells were analyzed for control condition measurements, and 639 segments from 11 cells for p190DN condition measurements.

## Acknowledgment

This work was supported by NIH grant R35GM136226 (L.H.), and Chan Zuckerberg Initiative (L.H.). L.H. is Irma T. Hirschl Career Scientist. We thank members of the Segall and Cox laboratories at Albert Einstein College of Medicine for their helpful discussions.

## Author contributions

S.P. and L.H. conceived the project. S.P. and L.H. designed the experiments. S.P. performed the experiments. S.P. and L.H. analyzed the results. S.P. and L.H. wrote and revised the manuscript.

---

**Supplemental Figure 1: RhoGDIgamma titration**

FRET/CFP emission ratios of the wild-type RhoB biosensor expressed alone or co-expressed with RhoGDIγ. A titration was performed by varying the RhoB biosensor–to– RhoGDIγ DNA ratio from 1:1 (200 ng of each plasmid) to 1:5 (200 ng RhoB biosensor plasmid and 1,000 ng RabGDI plasmid) in HEK-293T cells. Data are presented as mean ± SEM of 3 independent experiments. Statistical significance was assessed using an unpaired t-test. ns=non-significant, p > 0.05; *p < 0.05; **p<0.01; ***p < 0.001.

**Supplemental Figure 2: Activities of RhoB and TC10 biosensors at the cellular level (control and p190DN conditions) in MEFs constitutively expressing RHOB and TC10 biosensors**

**(A)** Whole-cell quantification of the FRET/donor ratio of the RhoB biosensor under control conditions and upon p190DN overexpression. Data are presented as mean ± SEM. Each dot represents a cell. 12 cells from 4 independent experiments for the control condition, and 11 cells from 3 independent experiments for the p190DN condition. Statistical significance was assessed using an unpaired t-test. ns=non-significant

**(B)** Whole-cell quantification of the FRET/donor ratio of the TC10 biosensor under control conditions and upon p190DN overexpression. Data are presented as mean ± SEM. Each dot represents a cell. 12 cells from 4 independent experiments for the control condition, and 11 cells from 3 independent experiments for the p190DN condition. Statistical significance was assessed using an unpaired t-test. ns=non-significant

**Supplemental Figure 3: Expression levels of RhoB and TC10, both endogenous proteins and biosensors, in MEFs constitutively expressing the RhoB and TC10 biosensors.**

**(A)** Quantification of the RhoB or TC10 biosensor–to–endogenous RhoB or TC10 ratio, respectively, by Western blot analysis. Each dot represents an individual sample. n=13.

**(B)** Quantification of the RhoB and TC10 endogenous protein with or without Dox. Each dot represents an individual sample. n=13.

**Supplementary Fig. 4: Temporal autocorrelation of the protrusion edge velocities**

Periodicity of cell edge protrusions was measured using autocorrelation functions from the morphodynamic analysis, in control (blue trace) and p190RhoGAP dominant-negative (p190DN)–transfected (red trace) MEFs constitutively expressing both RhoB and TC10 biosensors. Correlation curves were computed from n = 535 individual windows across 12 cells from 4 independent experiments for the control condition, and n = 639 individual windows across 11 cells from 3 independent experiments for the p190DN condition. Data are presented as mean ± 95% CI.

**Supplementary Movie 1**

CFP-YFP RHOB biosensor together with NIR-TC10 biosensor in MEFs cells, at 10s interval at 40x magnification in control condition. Frame playback rate: 7 fps, white bar = 10 μm.

**Supplementary Movie 2**

CFP-YFP RHOB biosensor together with NIR-TC10 biosensor in MEFs cells, at 10s interval at 40x magnification in p190DN condition. Frame playback rate: 7 fps, white bar = 10 μm.

