## Supplementary figures and images for "A new single chain, genetically encoded biosensor for RhoB GTPase based on FRET, useful for live-cell imaging"

### Supplementary Figures

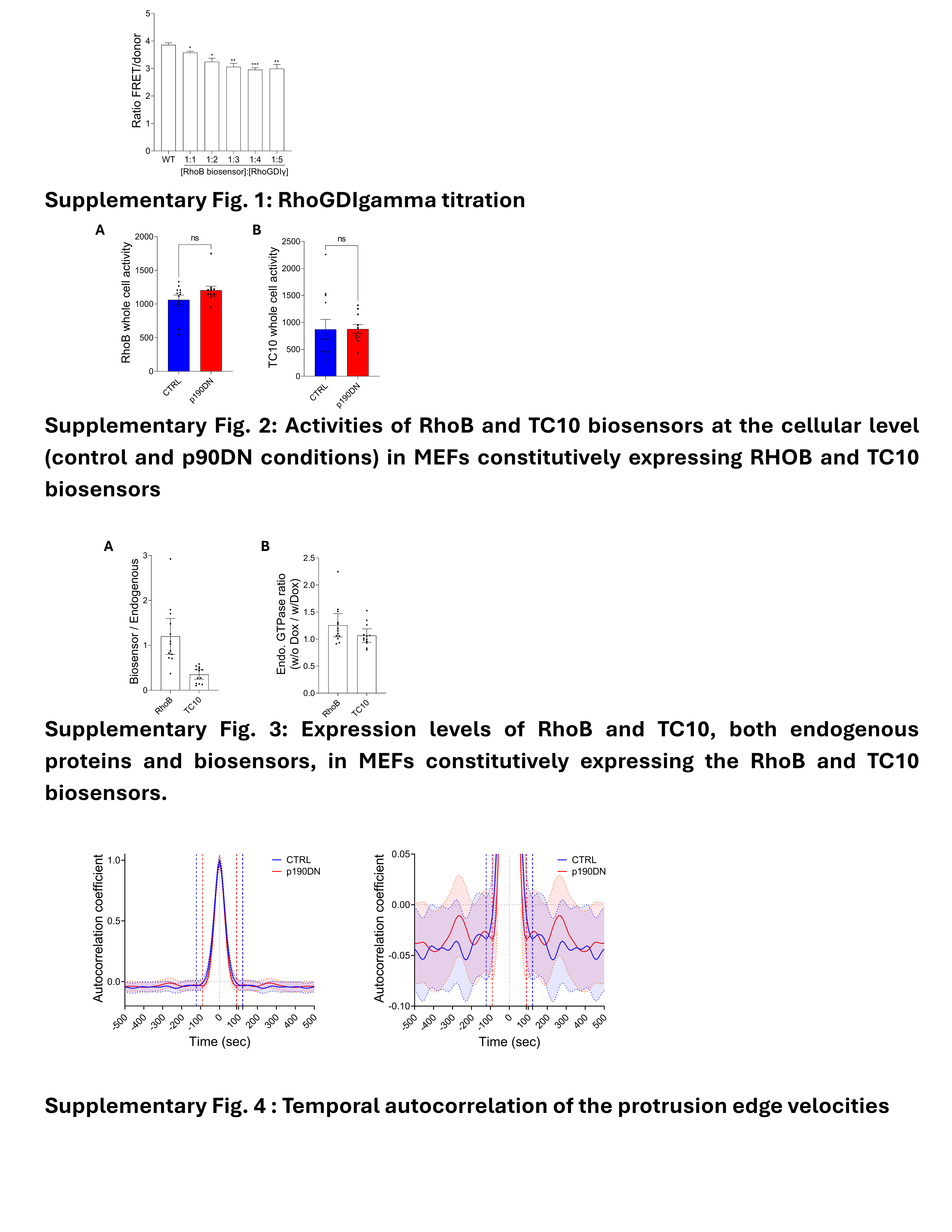
